# Functional Connectivity Evoked by Orofacial Tactile Perception of Velocity

**DOI:** 10.1101/843441

**Authors:** Yingying Wang, Fatima Sibaii, Rebecca Custead, Hyuntaek Oh, Steven M. Barlow

## Abstract

The cortical representation of orofacial pneumotactile stimulation involves a complex network, which is still unknown. This study aims to identify the characteristics of functional connectivity (FC) elicited by different saltatory velocities over the perioral and buccal surface of the lower face using functional magnetic resonance imaging (fMRI) in twenty neurotypical adults. Our results showed 25 cm/s evoked more functional coupling in the right hemisphere, suggesting 25 cm/s might be optimal velocity if bilateral brain damages occur. The decreased FC between the right secondary somatosensory cortex and right posterior parietal cortex for 5 cm/s versus All-on showed that the relatively slow velocity evoked less coupling in the ipsilateral hemisphere, which suggesting functional coupling in the contralateral hemisphere is in charge of orofacial tactile perception of velocity. The increased FC between the right thalamus and bilateral secondary somatosensory cortex for 65 cm/s versus All-on indicated that the neural encoding of relatively fast tactile velocity is more coupling between the right thalamus and bilateral secondary somatosensory cortex. Our results have shown different characteristics of FC for each seed at various velocity contrasts (5 > 25 cm/s, 5 > 65 cm/s, and 25 > 65 cm/s), suggesting the neuronal networks encoding the orofacial tactile perception of velocity. The difference of functional connectivity among three velocities may indicate the optimal stimulation setting for better therapeutic effects on stroke recovery.

## INTRODUCTION

The somatosensory system can process complex tactile stimuli from peripheral receptors through interactions between bottom-up thalamocortical and top-down corticocortical/cortico-thalamo-cortical pathways (Avivi-Arber et al., 2011;Lundblad et al., 2011;Zembrzycki et al., 2013;Rocchi et al., 2016;Hwang et al., 2017). The somatosensory system can process complex information about the location, velocity and transverse length of tactile stimuli, and has high cortical plasticity (Charlton, 2003). The cross-modality plasticity theory suggested that the somatosensory stimuli could evoke neural responses to promote learning new motor skills (Vidoni et al., 2010;Nasir et al., 2013;Ladda et al., 2014;Bernardi et al., 2015) and to perform accurate motor tasks (Pearson, 2000). Damages to the primary somatosensory cortex (SI) (e.g., by stroke, by traumatic brain injury, etc.) could result in orofacial sensory and motor deficits, the recovery of sensorimotor system requires changes in neuronal connections (Nudo, 2011;2013). Pneumotactile stimulation at different stimulus rates (2-6 Hz) on lower face have shown significant short- and long-term adaptation patterns in primary and secondary somatosensory cortices, and posterior parietal cortex (Popescu et al., 2013;Custead et al., 2015;Venkatesan et al., 2015). Thus, it is essential to study the changes of neuronal networks under certain somatosensory stimuli, which will benefit the development of optimal rehabilitation strategies for maximizing sensorimotor recovery in disease (Kaelin-Lang et al., 2002;Wu et al., 2006).

Previous functional magnetic resonance imaging (fMRI) studies have identified somatosensory networks including SI (subareas BA 3a, 3b, 1, 2), secondary somatosensory cortex (SII, BA40, 43), primary motor cortex (MI, BA 4), supplemental motor area (SMA, BA 6), posterior parietal cortex (PPC, BA 7), prefrontal cortex, and insular cortex (IC), as well as sensorimotor integration regions in the superior temporal gyrus (STG), supramarginal gyrus (SMG), thalamus, and cerebellum (Blatow et al., 2007;Huang et al., 2012). The sensory information like tactile motion perception on the face is received by the mechanoreceptors that project to the brain through the trigeminal nerve (Haggard and de Boer, 2014). The lack of neural bases of moving stimulation on face limited our understanding of velocity and directional encoding in the sensory domain. Our fMRI study is the first to identify a putative neural somatosensory velocity network with bilateral SI, bilateral cerebellum, bilateral middle occipital gyrus, left MI, right SII, right STG, and right SMG, right inferior frontal gyrus (IFG) (Custead et al., 2017). Brain regions elicited by the stimulus arrays demonstrated potential neurotherapeutic applications and rapidly adapting brain networks could be potentially used for monitoring or inducing brain plasticity and neural circuit reorganization after pneumotactile stimulation (Custead et al., 2017). Custead *et al.* have used univariate generalized linear model (GLM) that assumes the brain regions to be functionally specialized (sometimes termed ‘functional segregation’) rather than functionally integrated. However, this perspective limited our understanding of how different brain regions communicate with each other, which is important to understand complex neuronal networks (Tononi et al., 1998). Functional connectivity (FC) measured by correlation between time series of brain regions does not measure structural connections (e.g., axonal projections), but represents functional coupling between two or more spatially or anatomically distinct regions of the brain (Stevens, 2009).

The aim of the present study is to identify the characteristics of FC elicited by different saltatory velocities over the perioral and buccal surface of the lower face using our previous fMRI data (Custead et al., 2017). The pneumotactile simulator described in our previous work have activated SI, SII, and PPC using different velocities (Custead et al., 2017). Combining with literature (Blatow et al., 2007;Huang et al., 2012), ten regions of interests (ROIs) including bilateral SI, SII, PPC, dorsolateral prefrontal cortex, and thalamus were selected for ROI-to-ROI analysis. Four ROIs including bilateral SI and SII were chosen for Seed-to-Voxel analysis. We hypothesize that three velocities will evoke different FC in the brain. Our results may indicate the optimal stimulation setting for better therapeutic effects on recovery for diseases (e.g., stroke, traumatic brain injury, etc.) and provide insight into differences in the neurobiology of various therapeutic strategies (e.g., velocities, etc.).

## MATERIALS AND METHODS

### Participants

Twenty healthy, right-handed, native English-speaking adults (15 females), 18-30 years of age (mean ± SD: 22.3 ± 1.7), agreed to participate in the study after providing written informed consent. All participants had no history of neurological or psychiatric disorders, or any chronic illness or scheduled medications. The study was approved by the Institutional Review Board at the University of Nebraska-Lincoln.

### Paradigms

Our previous publication detailed the study design (Custead et al., 2017;Oh et al., 2017) (see supplementary figure 1). We used a block design and each twenty-second task block was followed by twenty-second resting block. The twenty-second block of 5 cm/s, or 25 cm/s, or 65 cm/s, or All-on, or All-off was randomized using a multichannel pneumatic amplifier and tactile array known as the Galileo Somatosensory™ system (Epic Medical Concepts & Innovations, Inc., Mission, Kansas USA). The Galileo uses probes known as chambered tactile cells (TAC-Cells) that are made from acetyl thermoplastic homopolymer and use tiny volumes of compressed air to rapidly deform the surface of the skin. The TAC-Cells are MRI-safe and incorporate a small capsule with a sealing flange, which can be adhered to the face using double-adhesive tape collars, with scalable and programmable control to create saltatory tactile arrays (see supplementary figure 1). The individual pressure pulses were transmitted through polyurethane tubing into the MRI suite, while the Galileo pneumatic amplifier and controller were located outside the MRI suite. The different velocities represented the different speeds of the air pressure pulses traveling (saltation) through channel 1 to 5 (see supplementary figure 1). For instance, the 5 cm/s indicated that the pressure pulses traveled through all channels sequentially within approximately 4 seconds. The 25 cm/s indicated that the pressure pulses traveled through all channels sequentially within approximately 1 second. The 65 cm/s indicated that the pressure pulses traveled through all channels sequentially within approximately 0.5 second. The All-on indicated that the pressure pulses traveled through all channels simultaneously. The All-off indicated that no pressure pulse traveled through all channels, which is equivalent to resting period.

### Stimulus device

Pneumotactile velocity stimuli were delivered to the facial skin by the Galileo system (see supplementary figure 1). For all stimuli, the Galileo system was programmed to generate biphasic pulses with duration of 60 ms, frequency of 1 Hz, 10 ms rise-fall time (10-90% intercepts), amplitude from −5 to 28 kPa. A laptop with windows 8.1 (64 bit) controlled the Galileo via an in-house software to generate sequential pressure pulses to channel 1 to 5. Pneumatic TAC-Cells were aligned on each participant from the right philtral column to the right (buccal) face. The individual array traverse length was calculated based on the distance between cells (each length measured from the center of one cell to center of the next). Because of bifurcation of the first two channels, both the upper and lower cells of those channels were considered ‘first’ and ‘second’ in the array. The measurement values of array length were used to designate on/off times for velocity sequences (traverse speed in cm/s). Therefore, velocity protocols were consistent across all participants, regardless of orofacial size. The resulting *.xml program produced a series of pneumotactile saltatory stimuli that traversed the skin in a repeating medial-to-lateral (upper/lower lips to lateral cheek) direction at three velocities (5, 25, 65 cm/s) as well as the ‘ALL-ON’ conditions. The stimulation array consisted of 7 small TAC-Cells that were adhered to the hairy skin of the right lower face, effectively sealing it to the skin. In this way, pressure dynamics within each cell resulted in skin deflection without acoustic or electrical artifact. Participants reported the resulting sensory experience as a moving sequence of discrete ‘taps’ or ‘raindrops’ on their lower face. These TAC-Cells are ported through a barb-fitting and connected to a 25 cm length of silicone tubing for flexible strain relief, and then coupled to a 5.18 m (1.6 mm in diameter) polyurethane line attached to the designated pneumatic ports on the GALILEO stimulus generator (see supplementary figure 1). The flanged surface of each TAC-Cell was secured to the skin using double-adhesive tape collars following skin preparation with tincture of benzoin to improve adhesion.

### Data acquisition

All images were acquired on a 3.0 T Siemens Skyra whole-body MRI system (Siemens Medical Solutions, Erlangen, Germany) with a 32-channel head coil. A high-resolution T1-weighted three-dimensional anatomical scan was acquired using magnetization-prepared rapid gradient-echo sequences (MPRAGE) with the following parameters: TR/TE/TA = 2.4 s/3.37 ms/5:35 minutes, flip angle = 7°, field of view = 256 × 256 mm, spatial resolution = 1 × 1 × 1 mm^3^, number of slices = 192. Following the MPRAGE anatomical scan, three sessions of functional MRI (fMRI) scans were recorded using a T2*-weighted echo planar imaging (EPI) sequence with the following parameters: TR/TE/TA = 2.5 s/30 ms/800 s, voxel size = 2.5 × 2.5 × 2.5 mm^3^, flip angle = 83°, number of slices = 41, number of volumes = 320.

Pneumotactile stimulus generation was synchronized to the MRI scanner using the first optical output TR TTL (transistor-transistor logic) pulse generator. The first TR pulse from the scanner at the onset of each fMRI acquisition was input to a Berkeley Nusleonics (Model 645) programmable pulse generator connected to the Galileo system. The pulse generator served as a timing mechanism sent triggers to the Galileo system to produce a velocity sequence every 40 seconds. The Galileo system generated a velocity condition for 20 seconds, then wait for the external trigger to initiate the next velocity sequence. In total, there were three sessions of the functional image acquisitions. Each session consisted of four sets of twenty-second block of any of the five conditions (5 cm/s, 25 cm/s, 65 cm/s, All-on, All-off were randomized) followed by twenty-second resting block. Each session consisted of 80-second 5 cm/s, 80-second 25 cm/s, 80-second 65 cm/s, 80-second All-on, and 80-second All-off, and 400-second resting period. In total, each session lasted approximately 800 seconds. Nineteen participants completed all three sessions, while one participant only completed two sessions.

### Data analysis

The CONN toolbox (Whitfield-Gabrieli and Nieto-Castanon, 2012) (http://www.nitrc.org/projects/conn) was used for pre-processing all images and compute brain connectivity using both seed-based and region-of-interests (ROIs)-based approaches. The CONN toolbox used Statistical Parametric Mapping (SPM12, http://www.fil.ion.ucl.uk/spm) to pre-process all image volumes including following steps: motion artefact corrections (realignment and scrubbing). Both toolboxes are based on MATLAB (MathWorks, Natick, U.S.A.). The functional data were realigned to correct for head motion, scrubbed for outliers, coregistered to the structural image and normalized to the Montreal Neurological Institute (MNI) space. Normalized images were smoothed with an isotropic Gaussian kernel of 8 mm FWHM.

The task-related functional connectivity was computed in the CONN toolbox. For each participant, CONN implemented CompCor to identify principal components associated with segmented white matter (WM) and cerebrospinal fluid (CSF) (Behzadi et al., 2007). These components were entered as confounds along with realignment parameters in a first-level analysis.

For the ROI-to-ROI analyses, we studied functional connectivity between ROIs for different velocities (e.g., 5 cm/s, 25 cm/s, 65 cm/s, All-on). A total of ten ROIs (five bilateral ROIs, see supplementary table 1) were created using MNI coordinates in the CONN toolbox and the MNI coordinates were based on the literature. The averaged fMRI time series were extracted from each ROI. The ROI-to-ROI correlation coefficients were obtained by calculating all possible correlation coefficients between the time series of each pair of ROIs.

For seed-based task-related functional connectivity analyses, we investigated connectivity between the four ROIs and all other voxels in the brain using seed-to-voxel analyses in the CONN toolbox. The four seeds (see supplementary table 1) including bilateral primary somatosensory cortex (SI) and secondary somatosensory cortex (SII) were chosen based on the literature to identify clusters of voxels in the brain exhibited the effects of velocity. The Seed-Voxel correlation coefficients were obtained by computing all possible cross-correlation coefficients between the time series of the seed and all residual voxels in the brain, and converting them to Z-scores. We used the second level analysis in CONN toolbox to compare functional connectivity among different velocity within group.

To control for multiple testing, a False Discovery Rate (FDR) correction of q < 0.05 was applied for all results (Benjamini and Hochberg, 1995).

## RESULTS

### ROI-based functional connectivity

In Figure 1, functional networks for each velocity (5 cm/s, 25 cm/s, and 65 cm/s) were overlaid onto three-dimensional rendered brain on the first row and task-related FC matrices for each velocity (5 cm/s, 25 cm/s, and 65 cm/s) were plotted on the second row (p < 0.05, FDR corrected). Comparing 5 cm/s and 25 cm/s task conditions, increased functional connectivity was identified between right DLPFC and right thalamus (see Figure 2). There is no significant difference of functional connectivity between 5 cm/s and 65 cm/s and between 25 cm/s and 65 cm/s for all ROI-to-ROI pairs. The contrast of 5 cm/s versus All-on condition showed significant decreased functional connectivity between right SII and right PPC and the contrast 65 cm/s versus All-on condition revealed increased functional connectivity between right thalamus and left SII and between right thalamus and right SII (see Figure 3).

**Figure 1.**
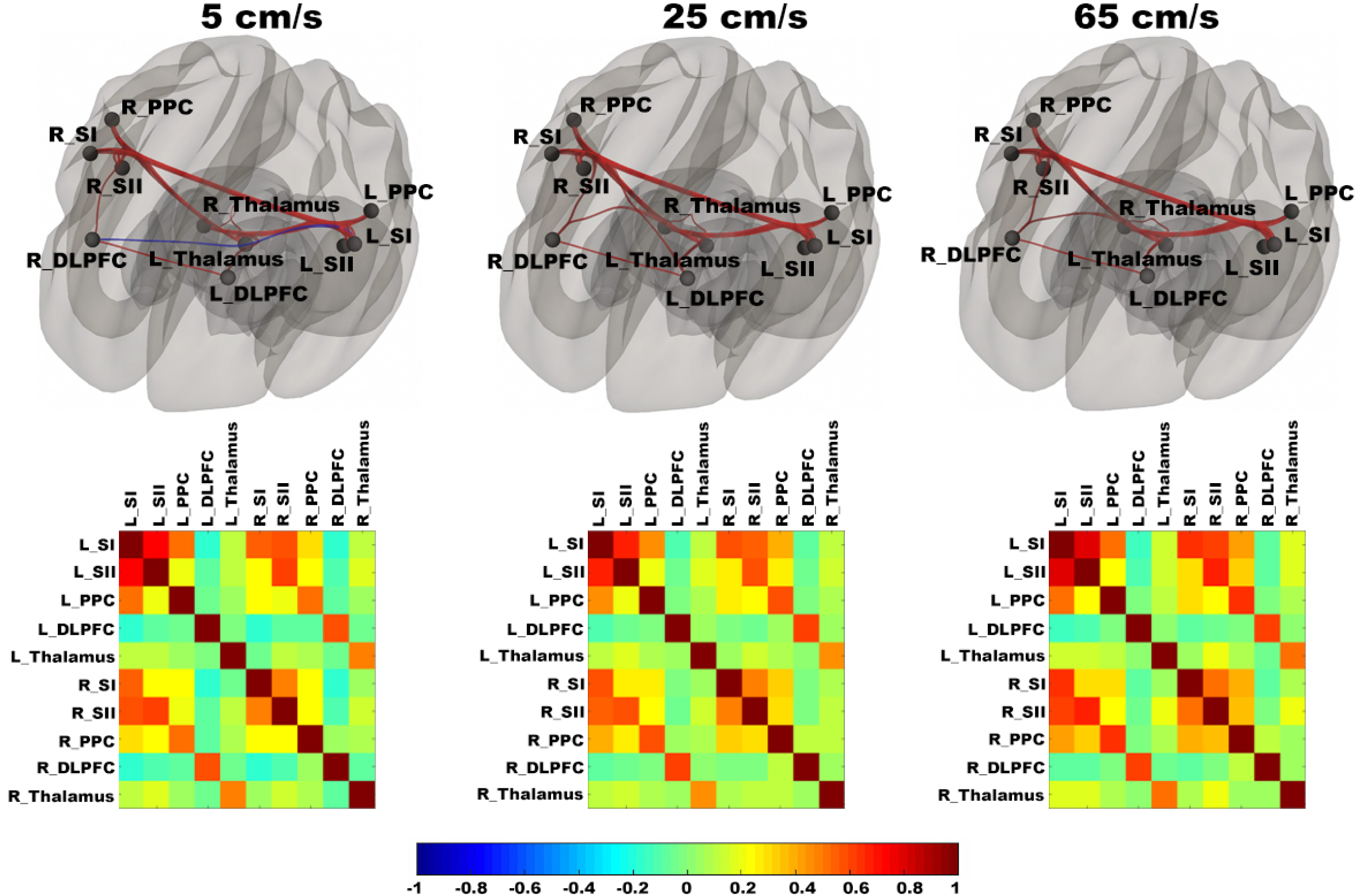
Shows ROI-to-ROI based connectivity maps (first raw) and connectivity adjacent matrices (second row) for three velocities (5, 25, 65 cm/s). Total six region of interests (ROIs) include bilateral primary somatosensory cortex (L_SI and R_SI), bilateral supplementary somatosensory cortex (L_SII and R_SII), bilateral Posterior Parietal Cortex (L_PPC and R_PPC), bilateral dorsolateral Prefrontal Cortex (L_DLPFC and R_DLPFC), and bilateral thalamus (L_Thalamus and R_Thalamus).

**Figure 2.**
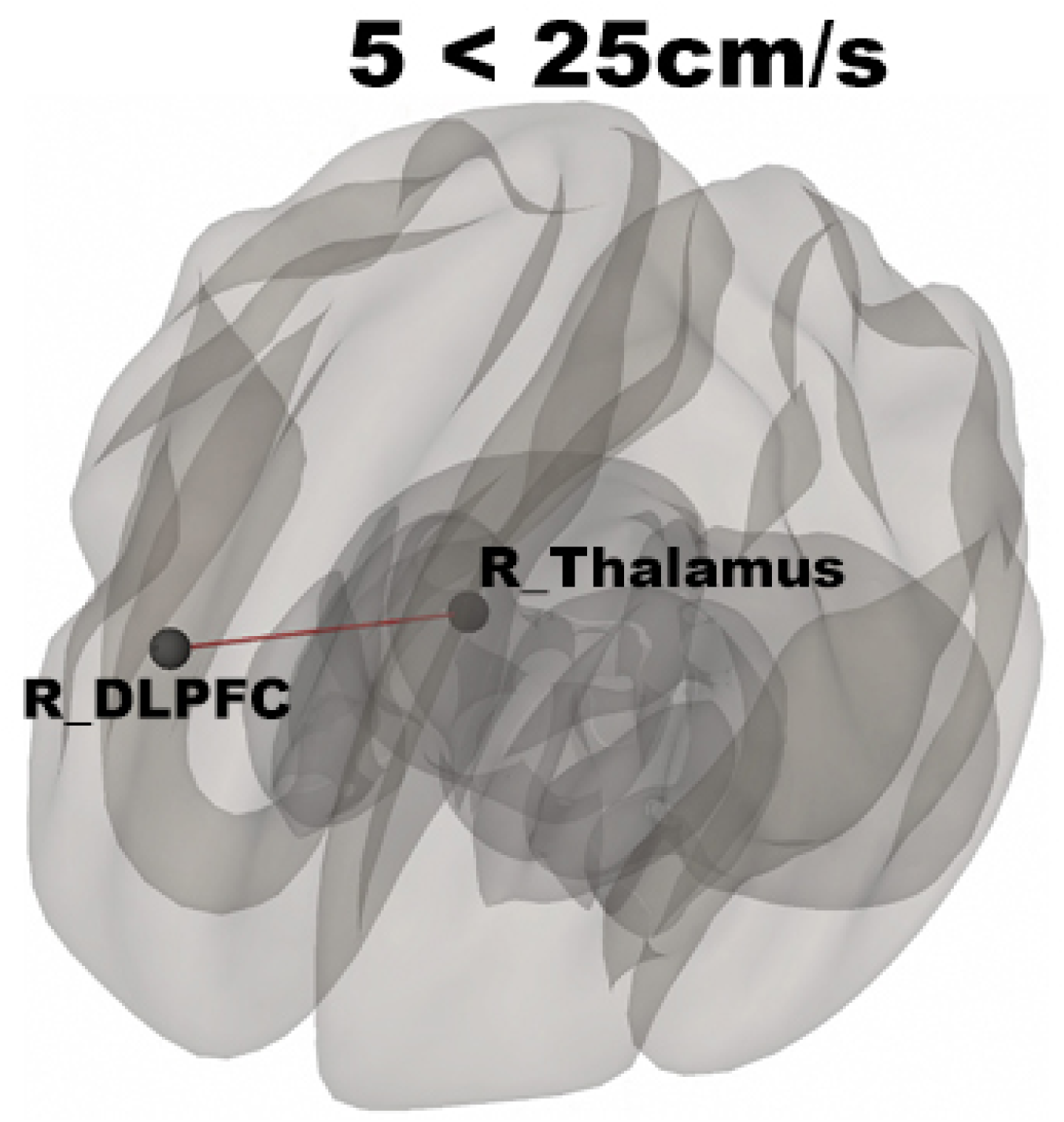
Shows increased connectivity between the right thalamus and right DLPFC for 25 > 5 cm/s contrast (p < 0.05, FDR corrected).

**Figure 3.**
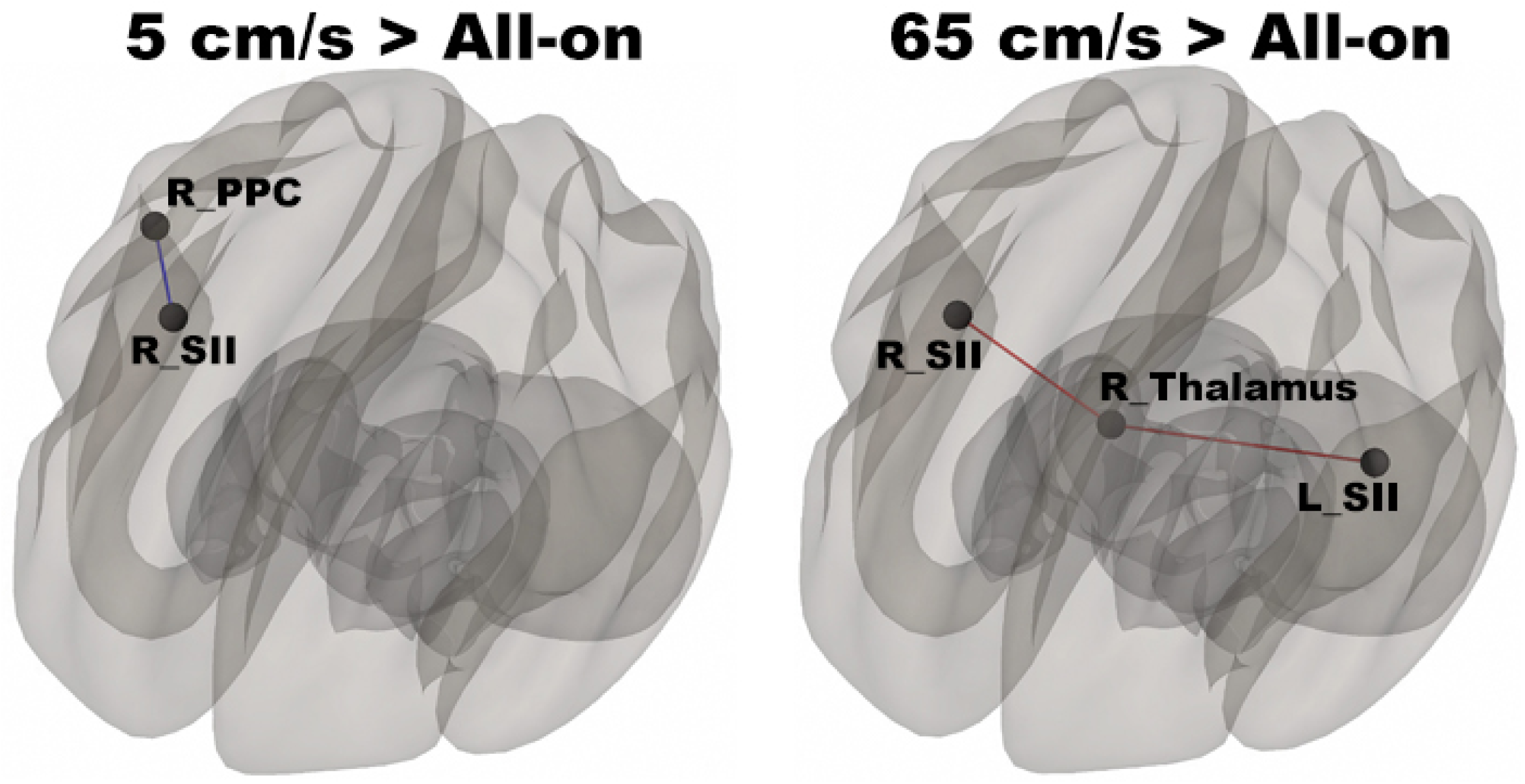
Shows increased connectivity between the right SII and the right PPC for 5 cm/s > All-on and increased connectivity between the right thalamus and the bilateral SII for 65 cm/s (p < 0.05, FDR corrected).

### Seed-based functional connectivity

The seed-to-voxel analysis assessed FCs between four seed regions covering bilateral SI and SII and all other voxels in the brain (p < 0.05, FDR corrected, cluster size > 35). For the left SI seed, results revealed increased FC in the left PostCG and left SMG for 5 cm/s > 25 cm/s (see Table 1 and Figure 4), in the right pMTG, right cerebellum 6, and right AG for 5 > 65 cm/s (see Table 1 and Figure 4), in the bilateral iLOC, right sLOC, right FG, right Cerebellum 6 for 25 > 65 cm/s (see Table 1 and Figure 4). For the left SII seed, increased FCs were only in the left SPL and right sLOC for 25 cm/s > 65 cm/s (see Table 1 and Figure 5). For the right SI seed, increased FCs were shown in the left iLOC and right pMTG along with decreased FCs in the right IC for 5 cm/s > 65 cm/s, and increased FCs were also observed in the bilateral iLOC along with decreased FC in the left cerebellum crus 2 for 25 > 65 cm/s (see Table 1 and Figure 6). For the right SII seed, decreased FC was present in the right SFG for both 5 > 65 cm/s and 25 > 65 cm/s. Additionally, increased FCs were shown in the left SPL and sLOC for 25 > 65 cm/s (see Table 1 and Figure 7).

**Table 1.** Seed-to-voxel results of changes in functional connectivity related to each velocity

|  |  |  |  |  |
| --- | --- | --- | --- | --- |
| Region | Coordinates in MNI space |  |  | Voxels |
|  | X (mm) | Y (mm) | Z (mm) |  |
| 1. Left SI Seed |  |  |  |  |
| Contrast: 5 > 25 cm/s |  |  |  |  |
| Left PostCG (35), aSMG (22) | -38 | -36 | 42 | 57 |
| Contrast: 5 > 65 cm/s |  |  |  |  |
| Right pMTG | 70 | -22 | -12 | 155 |
| Right Cerebellum 6 | 34 | -54 | -24 | 73 |
| Right AG | 60 | -52 | 26 | 64 |
| Contrast: 25 > 65 cm/s |  |  |  |  |
| Left iLOC (140) | -36 | -78 | 2 | 140 |
| Right iLOC (490), sLOC (138), FG (64) | 40 | -82 | -6 | 692 |
| Right FG (53), Right Cerebellum 6 (46) | 34 | -60 | -22 | 99 |
| 2. Left SII Seed |  |  |  |  |
| Contrast: 25 > 65 cm/s |  |  |  |  |
| Left SPL | -32 | -52 | 68 | 42 |
| Right sLOC | 36 | -84 | 18 | 42 |
| **3. Right SI Seed** |  |  |  |  |
| *Contrast: 5 > 65 cm/s* |  |  |  |  |
| Left iLOC | -42 | -72 | 2 | 78 |
| Right pMTG | 64 | -24 | -4 | 40 |
| *Contrast: 5 < 65 cm/s* |  |  |  |  |
| Right IC | 40 | -4 | -6 | 109 |
| *Contrast: 25 > 65 cm/s* |  |  |  |  |
| Left iLOC | -38 | -78 | 0 | 96 |
| Right iLOC | 46 | -82 | -4 | 298 |
| *Contrast: 25 < 65 cm/s* |  |  |  |  |
| Left Cerebellum Crus 2 | -38 | -66 | -54 | 34 |
| **4. Right SII Seed** |  |  |  |  |
| *Contrast: 5 < 65 cm/s* |  |  |  |  |
| Right SFG | 18 | 8 | 66 | 76 |
| *Contrast: 25 > 65 cm/s* |  |  |  |  |
| Left SPL | -30 | -58 | 62 | 147 |
| *Contrast: 25 < 65 cm/s* |  |  |  |  |
| Right SFG | 18 | 12 | 48 | 50 |
| Note. PostCG = Postcentral Gyrus, aSMG= anterior Supramarginal Gyrus, pMTG = posterior Middle Temporal Gyrus, AG = Angular Gyrus, iLOC = inferior Lateral Occipital Cortex, sLOC = superior Lateral Occipital Cortex, FG = Fusiform Gyrus, SPL = Superior Parietal Lobule, IC = Insular Cortex, SFG = Superior Frontal Gyrus. |  |  |  |  |

**Figure 4.**
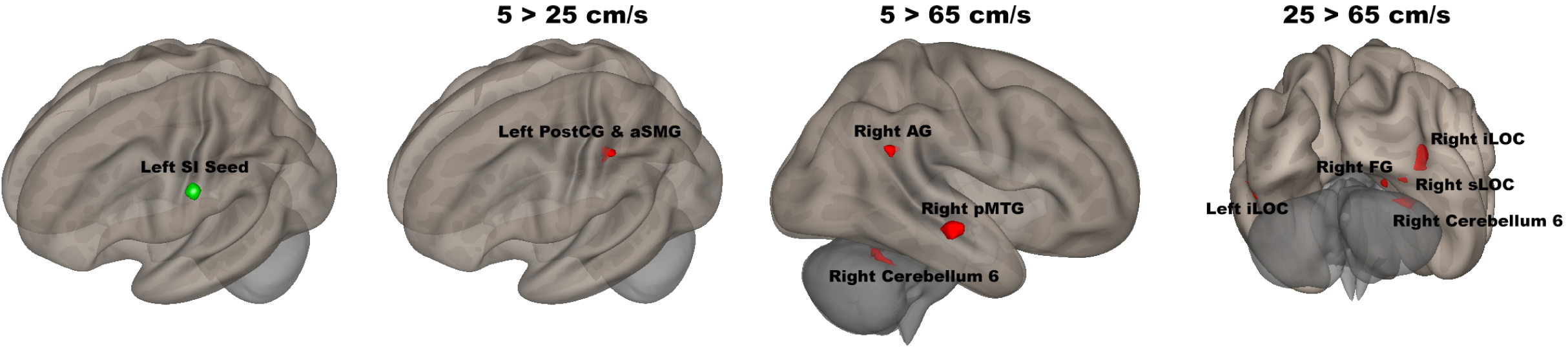
Shows the left SI seed (green sphere) overlaid on a standardized three-dimensional template and the seed-to-voxel results were presented on the right (p < 0.05, FDR corrected).

**Figure 5.**
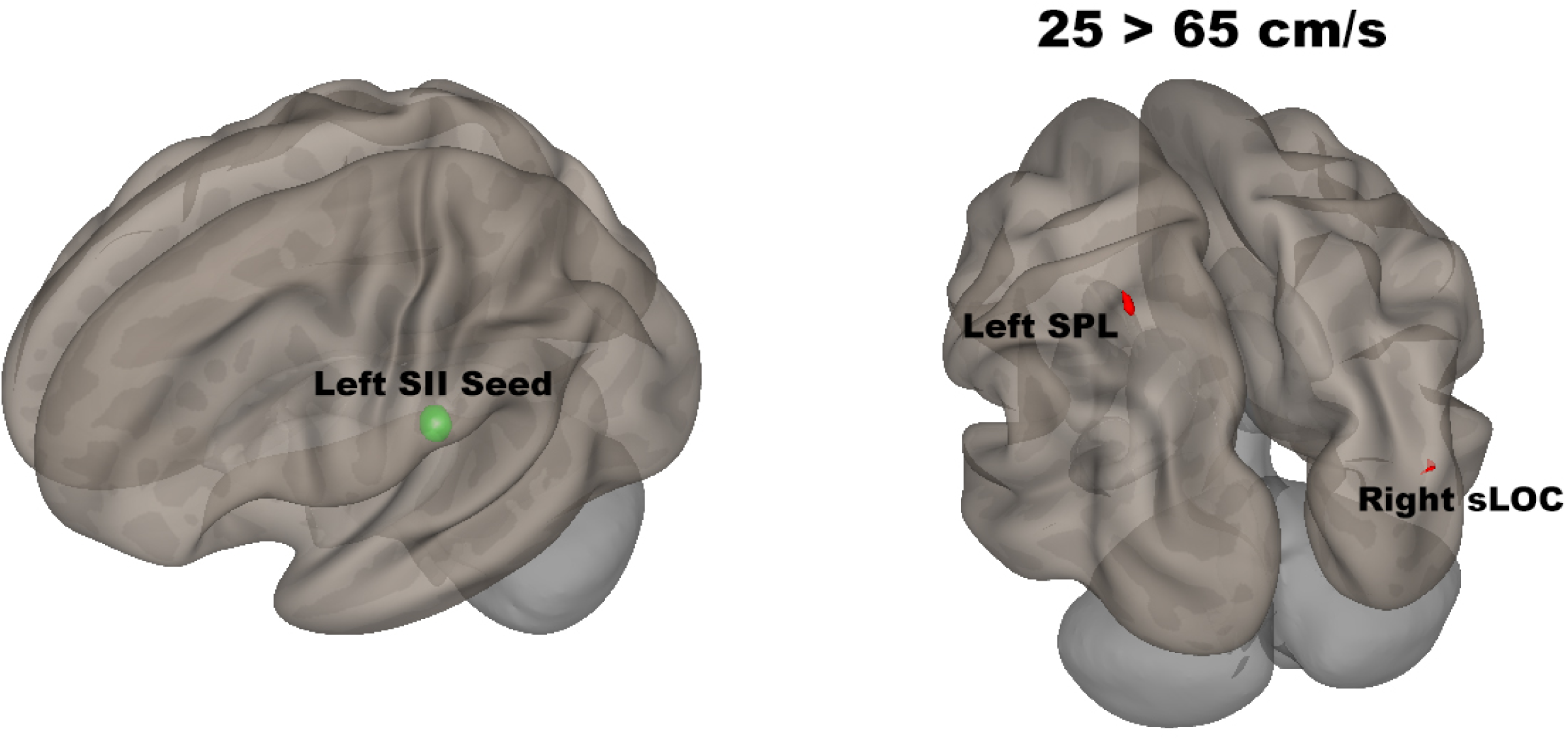
Shows the left SII seed (green sphere) overlaid on a standardized three-dimensional template and the seed-to-voxel results were presented on the right (p < 0.05, FDR corrected).

**Figure 6.**
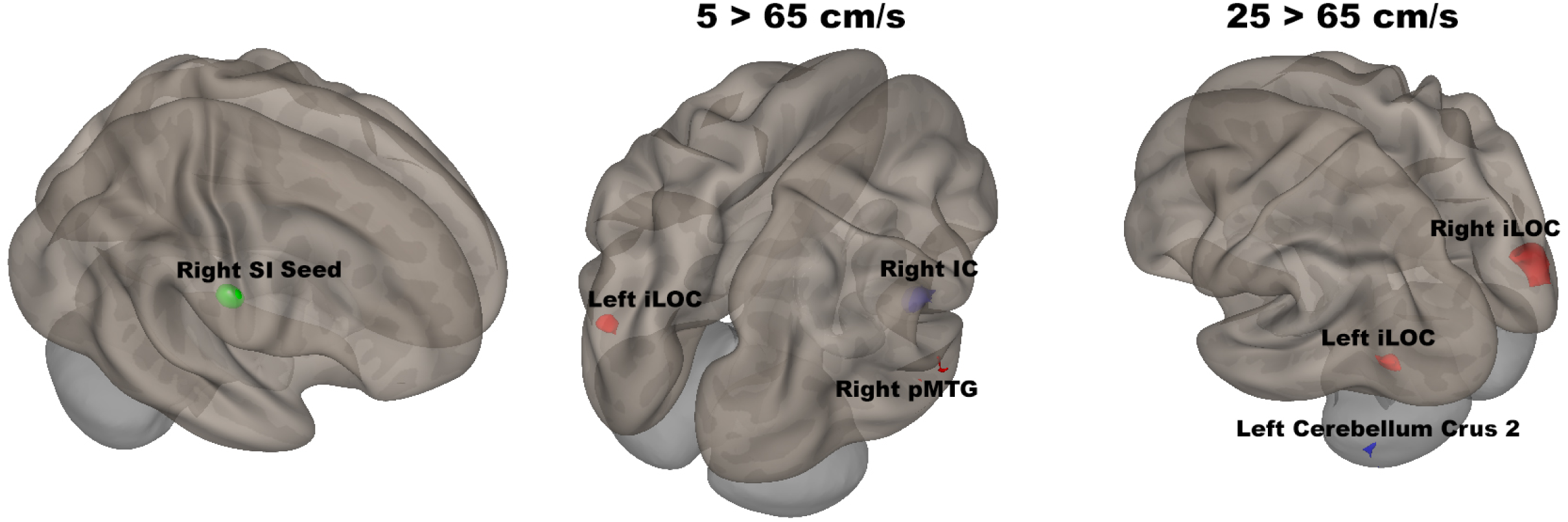
Shows the right SI seed (green sphere) overlaid on a standardized three-dimensional template and the seed-to-voxel results were presented on the right (p < 0.05, FDR corrected).

**Figure 7.**
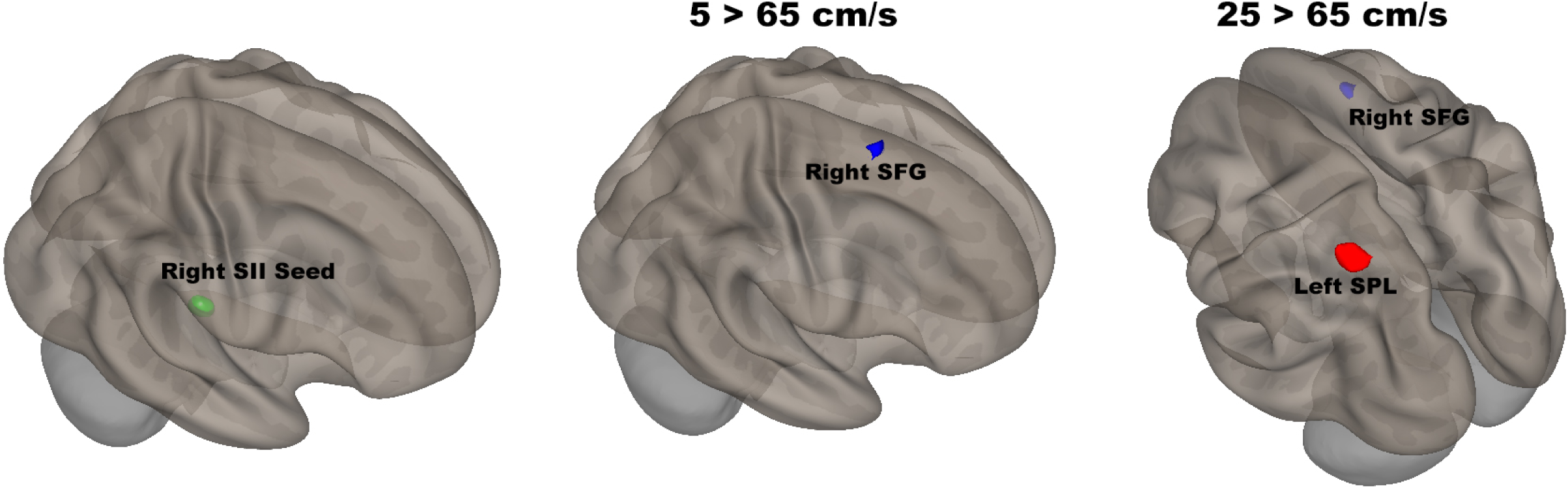
Shows the right SII seed (green sphere) overlaid on a standardized three-dimensional template and the seed-to-voxel results were presented on the right (p < 0.05, FDR corrected).

## DISCUSSION

The present study examined FC evoked by the orofacial tactile perception of velocity using fMRI in 20 neurotypical adults. This is the first attempt to identify FC evoked by novel saltatory pneumotactile stimuli using TAC-Cells with the Galileo system. Our ROI-to-ROI results showed 25 cm/s evoked more functional coupling in the right hemisphere (ipsilateral to the tactile stimuli) (see Figure 1), suggesting 25 cm/s might be optimal velocity if bilateral brain damages occur. The decreased FC between the right SII and right PPC for 5 cm/s versus All-on showed that the relatively slow velocity evoked less coupling in the ipsilateral hemisphere, which suggesting functional coupling in the contralateral hemisphere is in charge of orofacial tactile perception of velocity. The increased FC between the right thalamus and bilateral SII for 65 cm/s versus All-on indicated that the neural encoding of relatively fast tactile velocity is more coupling between the right thalamus and bilateral SII. The Seed-to-Voxel approach used bilateral SI and SII seeds to identify different network patterns for each velocity. Our results have shown different characteristics of FC for each seed at various velocity contrasts (5 > 25 cm/s, 5 > 65 cm/s, and 25 > 65 cm/s), suggesting the neuronal networks encoding the orofacial tactile perception of velocity.

### Orofacial Tactile Perception of Velocity

Our paradigm passively delivered the tactile stimuli with various velocities (5, 25, 65 cm/s) to the right side of participants’ face. For the All-on condition, the Galileo system delivered pressure pulses to all five channels simultaneously. The contrasts of velocity versus All-on condition (5, 25, or 65 > All-on) revealed FC evoked by the orofacial tactile perception of velocity. For 5 cm/s > All-on, reduced FC between the right SII and right PPC suggested less coupling in the right (ipsilateral) hemisphere, which aligns with our previous GLM results (Custead et al., 2017). The GLM results showed bilateral activation patterns when comparing 5 cm/s versus All-on. However, the GLM results are limited to the strength of the blood oxygen level depend (BOLD) signals and cannot determine the communication between brain regions, while FC analysis allows us to understand the coupling between brain regions. FMRI study has reported the representations of six body parts (face, fingers, legs, shoulders, lips, and toes) in the superior PPC (Huang et al., 2012). The increased FC between the right thalamus and bilateral SII for 65 cm/s > All-on indicated the role of thalamus as an integrative hub for functional brain networks (Hwang et al., 2017). Early animal study found that the SII receives substantial inputs from topographically appropriate regions within the ipsilateral ventrobasal nucleus and from the ipsilateral posterior group (Carvell and Simons, 1987), which proposed that SII in mice may complement the function of SI by helping to define the overall sensory context in which detailed tactile discriminations are made. Therefore, our results of the relatively faster velocity stimuli evoked stronger couplings between the right thalamus and bilateral SII suggested that SII in human may play an important role of discriminating velocity of orofacial tactile stimuli (Carvell and Simons, 1987;Tommerdahl et al., 2005a;Tommerdahl et al., 2005b).

For contrast 5 >25 cm/s, the ROI-to-ROI results showed significantly decreased FC between the right thalamus and the right DLPFC, which might be due to the increase of velocity requiring high level cognition (e.g., right DLPFC for attention and executive function) to decipher the tactile stimuli. The changes of temporal density of pneumotactile stimulation might drive the changes of neuronal populations in the brain during the orofacial tactile stimuli. The Seed-to-Voxel results indicated that the increased FC between the left SI seed and left PostCG/aSMG. Our previous fMRI findings have shown that the lowest temporal density of pneumotactile stimulation (5 cm/s) evoked the largest spatial extent of bilateral brain activity (Custead et al., 2017). Our FC results are complementary to our previous GLM results. Comparing to 25 cm/s, the low velocity 5 cm/s evoked stronger FC in the contralateral hemisphere but weaker FC in the ipsilateral hemisphere. Thus, the FC evoked by the low velocity 5 cm/s is stronger than the FC evoked by the mid-range velocity 25 cm/s in the contralateral hemisphere, align with other studies in the literature (Dreyer et al., 1979;Lamb, 1983;Whitsel et al., 1986).

For contrast 5 > 65 cm/s, we observed increased FCs in the left SI seed between the right AG, right pMTG, and right cerebellum 6, and increased FCs between the right SI seed and left iLOC and right pMTG, as well as decreased FCs between the right SI seed and right IC, and between the right SII seed and right SFG. The relatively slow velocity (5 cm/s) evoked stronger coupling between the left SI and left PostCG/left aSMG compared to the mid-range velocity (25 cm/s). The moving tactile stimuli presented at a low velocity such as 5 cm/s can appear to be processed in neuronal networks as discrete stimuli rather than a constant motion across the skin (Dépeault et al., 2013), which may contribute to the differences of FC versus other velocities. With the increases of stimulus velocity, enough loss of temporal and spatial details may lead to reduced discrimination accuracy (Lamb, 1983). The right AG, right pMTG, right cerebellum regions, and right IC regions have corresponded to our previous GLM results (Custead et al., 2017). Both MTG, SMG, and IC have been reported to be part of a ventral attention network responsible for bottom-up attention and sensorimotor response inhibition (Corbetta et al., 2008;Igelström and Graziano, 2017). Moreover, the cerebellar involvement is consistent with the putative role of the cerebellum in feedforward control of sensory-guided movements at 5 cm/s (Custead et al., 2017).

For contrast 25 > 65 cm/s, there were increased FCs between the left SII seed and left SPL/right sLOC, the increased FCs between the right SI seed and bilateral iLOC and between the right SII seed and left SPL. The interhemispheric increased FCs We also observed the decreased FC between the right SI seed and the left cerebellum crus 2, and between the right SII seed and the right SFG. Other studies have suggested the optimal range for accurate discrimination of tactile velocity is between 3 and 30 cm/s (Dreyer et al., 1979;Lamb, 1983;Whitsel et al., 1986). At higher velocity like 65 cm/s, participants are still able to discriminate the moving stimuli but with lower accuracy (Lamb, 1983). Therefore, the differences of FC patterns were presented between the two velocities.

The changes of FC for different velocities suggested that all three velocities can be used to induce neural plasticity and changes in neuronal connections. But each velocity has its uniqueness and can be used based on the sensitivity and spatial specificity needed for the specific neurotherapeutic applications.

### Contralateral versus Ipsilateral hemisphere

Animal studies have already reported both contralateral and ipsilateral activation during unilateral or bilateral activation (Tommerdahl et al., 2010). Our results support the view of an ipsilateral influence on SII, align with other studies (Tommerdahl et al., 2005a;Tommerdahl et al., 2006). Neurons in SII most often have bilateral receptive fields, unlike neurons in SI. (Whitsel et al., 1969).The present study depicted that reduced FC between the right PPC and right SII for 5 cm/s > All-on, suggesting the contralateral hemisphere evoked by the slow velocity is critical for neuronal encoding of orofacial tactile perception of velocity. We observed changes of FC in both hemispheres, in align with our previous report on bilateral cortical responses (Custead et al., 2017). The different velocities evoked different brain connectivity patterns. The interhemispheric (callosal) connections indicated pneumotactile stimulation reach SI via a two-stage pathway involving interhemispheric (callosal) connections between information processing levels higher than SI and subsequently via interhemispheric (corticocortical) projections to the SI face region. Our results are also align with that the human somatosensory system processes the tactile stimuli in a hierarchical scheme of somatosensory processing (Inui et al., 2004;Tommerdahl et al., 2010).

### Limitations

The present study has several limitations. First, the major limitation is the imaging modality that measures relatively slow hemodynamic responses on the order of second. FMRI data can provide some indirect measures to decode how the sensory system perceives different stimuli with various velocities. However, humans can make sensory decisions in less than 200 ms (Thorpe et al., 1996), which relies primarily on rapid synaptic neurotransmission on a time scale of millisecond (Kohn et al., 2002;Kohn and Whitsel, 2002). Thus, electrophysiology-based imaging approaches (i.e., magnetoencephalography, electroencephalography) are more suitable to study the dynamic changes of this rapidly changing system (Puts et al., 2019). Second, the relatively small sample size and wider age range can limit the power of this study. Lastly, no behavioral measures are collected to check the individual differences of perception ability.

### Conclusions and Future Directions

The present study found a distinct cortical connectivity pattern associated with each velocity. Our results demonstrated that the left SI evoked more connections than the left SII at three velocity settings (5 cm/s, 25 cm/s, 65 cm/s). The brain circuits changing with pneumotactile stimulation at different velocity settings indicate brain process tactile stimulation with different velocity settings differently. Physiologically, suprathreshold mechanical touch signals start as widespread, relatively diffuse activity across somatosensory macrocolumns that are driven by the characteristics of the stimulus. The paradigm in this study modulates neural circuits through changes in the velocity of a stimulus over a set block of time. While the velocity changes, neuronal populations may be driven by changes in the temporal density of pneumotactile stimulation. Animal and human studies have shown passively evoked sensory stimulation can enhance neuronal activity after stroke (Whitaker et al., 2007). Therefore, the present study has implications for applying various velocities to orofacial stimuli in order to bolster recovery for sensorimotor rehabilitation. For instance, combined with physical therapy for stroke patients or brain-injured survivors, it might induce more brain plasticity during rehabilitation (Small et al., 2002;Luft et al., 2005;da Guarda and Conforto, 2014).

## Contribution to the field

This is the first study using functional magnetic resonance imaging (fMRI) to identify the characteristics of functional connectivity elicited by novel pneumotactile stimulations with different velocity on the lower face. Our findings will elucidate the neuronal networks encoding the orofacial tactile perception of velocity. The difference in functional connectivity among three velocities may indicate the optimal stimulation setting for better therapeutic effects on stroke recovery.

## Ethics statements

### Studies involving animal subjects

Generated Statement: No animal studies are presented in this manuscript.

### Studies involving human subjects

Generated Statement: The studies involving human participants were reviewed and approved by University of Nebraska-Lincoln. The patients/participants provided their written informed consent to participate in this study.

### Inclusion of identifiable human data

Generated Statement: No potentially identifiable human images or data is presented in this study.

## Data availability statement

Generated Statement: The datasets generated for this study are available on request to the corresponding author.

## Acknowledgments

We thank the families for their participation. We also thank the supports for undergraduate research assistants from the UNL UCARE program funded in part by gifts from the Pepsi Quasi Endowment and Union Bank & Trust.

## Ethics Statement

The study was approved by the Institutional Review Board at University of Nebraska-Lincoln. Written informed consent was obtained from each participant in accordance with the Declaration of Helsinki.

## Author Contributions

YW proposed and performed connectivity analysis, and drafted the manuscript. SM contributed to the conception, design, and data collection of the study, and revising the manuscript critically for important intellectual content. RC and HO carried out the experiment and data collection. FS organized and pre-processed the data. All authors read and approved the submitted version and agree to be accountable for all aspects of the work.

## Conflict of Interest

The authors declare that the research was conducted in the absence of any commercial or financial relationships that could be construed as a potential conflict of interest.

## Funding

Thanks for funds from the Barkley Trust, Nebraska Tobacco Settlement Biomedical Research Development, College of Education and Human Sciences, and the Office of Research and Economic Development.

## Data Availability Statement

The raw data supporting the conclusions of this manuscript will be made available by the authors, without undue reservation, to any qualified researcher.

## Supplementary Material

The Supplementary Material for this article can be found online.

