## Supplements for "Functional Connectivity Evoked by Orofacial Tactile Perception of Velocity": Data Sheet 1.PDF

### *Supplementary Material*

#### Supplementary Tables and Figures

**Supplementary Table 1. Region of Interests**

| Name | Description | Coordinates in MNI space | | |
|---|---|---|---|---|
| | | X (mm) | Y (mm) | Z (mm) |
| Left Hemisphere | | | | |
| Left SI | ROI & Seed | -55 | -19 | 24 |
| Left SII | ROI & Seed | -48 | -24 | 16 |
| Left PPC | ROI | -56 | -31 | 32 |
| Left DLPFC | ROI | -27 | 32 | 36 |
| Left Thalamus | ROI | -9 | -17 | 6 |
| Right Hemisphere | | | | |
| Right SI | ROI & Seed | 56 | -13 | 29 |
| Right SII | ROI & Seed | 48 | -24 | 16 |
| Right PPC | ROI | 56 | -31 | 32 |
| Right DLPFC | ROI | 30 | 35 | 34 |
| Right Thalamus | ROI | 10 | -19 | 6 |

*Note:* MNI = Montreal Neurological Institute; ROI = region of interest; SI = primary somatosensory cortex, SII = supplementary somatosensory cortex; PPC = Posterior Parietal Cortex; DLPFC = dorsolateral Prefrontal Cortex

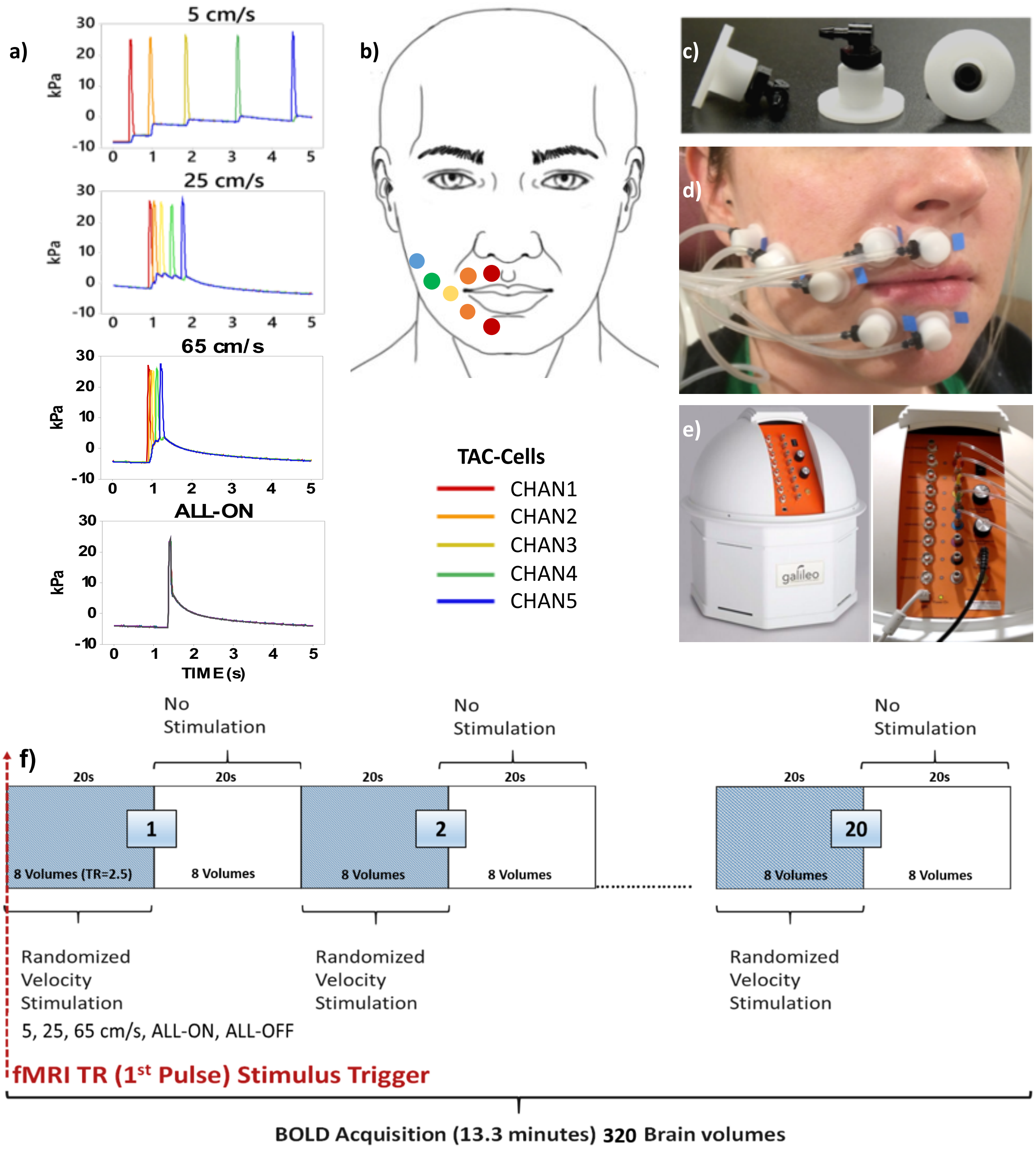


**Supplementary Figure 1.** Shows the experimental configuration for the Galileo somatosensory stimulator with pneumatic velocity array and experimental design. a) programmed time delays between pressure pulses at each cell resulted in 4 task conditions: 5 cm/s, 25 cm/s, 65 cm/s, ALL-ON synchronous activation (1 Hz), b) pneumatic cells were aligned on the participant from the right philtral column to the right buccal face, c) TAC-Cells: white flanged surface was adhered to skin surface with double adhesive colars, d) a series of pneumotactile saltatory stimuli traversed the skin in a repeating medial-to-lateral direction at 4 task conditions, e) the Galileo somatosensory simulator, f) mixed design for the study paradigm.
